# NEURA: A proof-carrying framework for hallucination-resistant neuroimaging automation

**DOI:** 10.64898/2026.04.27.721217

**Authors:** Jun Xie, Jing Wang, Xiumei Wu, Xinyuan Liu, Yiqi Mi, Qingjin Liu, Tong Xu, Chen Liu, Huafu Chen, Jing Guo

## Abstract

Neuroimaging research depends on heterogeneous software, multimodal data and multistage statistical workflows. Large language model (LLM)-based agents offer a route to automate these workflows, but their susceptibility to hallucination limits their credibility in scientific use. Here we introduce NEURA, a proof-carrying framework for hallucination-resistant neuroimaging automation. NEURA converts free-text research questions and neuroimaging datasets into executable analysis plans, validated outputs and structured reports. The system combines disease- and tool-aware planning with a deterministic verification layer inspired by formal proof: before any claim is retained for reporting, it must be checked against tool-derived evidence and domain axioms. On NeuroEval, an expert-curated benchmark of 110 neuroimaging tasks, NEURA achieved 89.5% planning accuracy, a 30.5% improvement over direct LLM queries. In a controlled hallucination-injection experiment, the verification layer detected all the injected error classes under the specified axiom bank and trust assumptions, with no false positives. In case studies of spinocerebellar ataxia type 3, NEURA reproduced cerebellar atrophy and abnormal diffusion patterns consistent with established pathology and independent expert analyses. Together, these findings show that coupling domain-grounded agency with proof-carrying verification can turn LLM-driven workflow automation from probabilistic self-checking into auditable scientific computation.

## Introduction

Neuroimaging is now widely recognized as a core experimental paradigm for investigating the structural and functional architecture of the human brain^1^, providing quantitative, noninvasive biomarkers for diagnosis, prognosis, treatment evaluation, and research on pathological pathways in neurological and psychiatric disorders^2,3^. As the field moves towards population-scale cohorts, higher-resolution acquisition and multimodal integration across structural, functional and diffusion magnetic resonance imaging (MRI) and large initiatives such as the UK Biobank^4^, ABCD^5^ and ENIGMA^6^ have created unprecedented opportunities to link brain imaging with genetics, development, ageing, disease heterogeneity and longitudinal clinical phenotypes. These resources have substantial clinical value, but their analysis remains challenging^7^. However, even with mature toolchains^8^ such as SPM^9^, FSL^10,11^, FreeSurfer^12^, and AFNI^13^ and workflow engines such as Nipype^14^, researchers must coordinate heterogeneous software ecosystems, file formats, parameters and intermediate outputs. All of these depend on expertise that spans neuroscience, data processing and software engineering, limiting reproducibility, scalability and clinical translation^15^. These challenges highlight the urgent need for analysis systems that can convert large-scale imaging resources into traceable and biologically meaningful evidence—an area where recent advances in multiagent systems offer transformative potential^16,17^.

The development of artificial intelligence (AI) systems for neuroimaging research presents several inherent challenges: (1) multidisciplinary reasoning and workflow composition (effective analyses must map disease concepts onto relevant brain regions, imaging modalities, preprocessing strategies, statistical models and reporting conventions^8^), (2) normative execution control (neuroimaging workflows are governed by strict methodological and quality-control standards, whereas open-loop agents often lack explicit structural constraints and fine-grained state governance^18^), (3) long-horizon state management (workflows that span dozens or hundreds of actions can suffer domain hallucination^19^, context loss and state drift^20^ when long-term memory and goal tracking are weak), (4) continual knowledge updating and tool extensibility (both neuroimaging evidence and the software ecosystem evolve rapidly^8,16^) and (5) transparency and traceability (scientific deployment requires explicit, auditable rationales that support trust, reproducibility and accountability^17^).

Concurrently, large language models (LLMs) have rapidly evolved into general-purpose foundation models that are reshaping scientific computing^21^. Beyond natural-language generation, recent evidence highlights their emergent capacity for algorithmic reasoning and complex problem decomposition ^17,22,23^. This evolution has catalysed the rise of “agentic AI”, which is a paradigm that has been successfully deployed in software engineering^24^ and synergizes LLM-based reasoning with the execution of external tools. This synergy enables systems to plan, act, see what happens, and then improve their methods over time. In scientific and biomedical domains, recent systems have shown that tool-augmented agents can support autonomous chemical experimentation, chemical reasoning, medical workflow automation and traceable rare-disease diagnosis^16,17,25^. However, a fundamental tension persists: LLM agents can generate domain-specific hallucinations that are syntactically fluent yet violate established constraints, and existing mitigation strategies that rely on LLM-based self-evaluations create “LLM-judges-LLM” circularity^19,26^ that cannot meet the demands of deterministic guarantees foe scientific applications.

Here, we present NEURA, which is an LLM-powered agentic system driven by a cognitively inspired architecture that pairs flexible reasoning with formal verification. NEURA uses specialized LLM modules (Planner, Executor, Reviewers) to autonomously plan and execute tools across heterogeneous imaging modalities, while a proof kernel provides deterministic claim-level verification independent of any LLM.

The system stands out because of four main architectural innovations that ensure that NEURA is strict and can be expanded. First, a dual knowledge graph framework grounds automated reasoning: a disease–region knowledge graph (DiseaseKG) encodes pathological associations between neurological conditions and brain structures, whereas a tool knowledge graph (ToolKG) maps functional dependencies, input/output formats, and usage constraints. Both graphs evolve through literature retrieval and execution feedback. Second, inspired by the concept of “skill”^27^ learning, we developed an open and extensible tool interface that integrates heterogeneous analysis methodologies and multimodal data formats. This architecture empowers the agent to autonomously distill reusable analytic skills from the methodological literature, thereby dynamically expanding its procedural repertoire to adapt to novel workflows and complex multimodal paradigms without manual reconfiguration. Third, a mixture-of-experts review (MoER) mechanism routes intermediate outputs to domain-specific reviewers that evaluate against expert-curated normative specifications, providing soft guidance on parameter legality, statistical conventions, and methodological quality at every workflow stage.Fourth, a Curry-Howard-inspired proof kernel^28^ provides a deterministic verification layer that operates independently of any LLM. Every claim in the generated report must carry a closed proof term that type-checks against a bank of neuroimaging axioms covering disease-region mappings, tool modality constraints, statistical prerequisites, and pipeline dependencies. Claims without valid proofs are automatically stripped, ensuring correctness by construction rather than LLM evaluation. Together with a memory-augmented reflection loop and self-healing code generation, this design maintains long-horizon consistency and supports recovery from runtime failures^18,20,29^.

We also established NeuroEval, an expert-curated benchmark that comprises 110 literature-derived neuroimaging tasks that span multiple disease domains and two levels of workflow complexity. On this benchmark, NEURA achieved 89.5% planning accuracy and substantially outperformed direct LLM queries, with average gains of 30.5% in planning accuracy, 25.6% in tool selection and 36.7% in tool ordering. In a controlled hallucination-injection experiment (9 types, 990 corrupted reports), the proof kernel reached F1=1.0 with zero false positives, whereas a DeepSeek-V3.2 LLM-as-judge baseline reached only F1=0.672 and completely failed on subtle methodological violations. In clinical case studies of spinocerebellar ataxia type 3 (SCA3), NEURA autonomously completed structural and diffusion MRI analyses and identified cerebellar atrophy and abnormal diffusivity patterns consistent with established pathology and independent expert manual analyses.

## 2. Results

### 2.1 Overview of NEURA

NEURA is an LLM-powered agent designed to reduce the complexity of neuroimaging analysis. It uses a structure that mimics how human researchers think, reason, and learn. In traditional workflows, neuroimaging analysis requires a deep accumulation of multidisciplinary knowledge. This requirement makes it challenging. NEURA fills the most important gap between theoretical scientific research and practical application.

The overall workflow of NEURA is illustrated in Fig. 1 and comprises three principal phases—understanding and planning, execution, and validation—preceded by an environment-aware parsing step that grounds the entire process in the actual data workspace. Given a research task defined by a natural-language query and a local workspace that contains neuroimaging datasets, NEURA first inspects the computational environment to identify available data modalities, file organization and metadata. This step grounds subsequent reasoning in the actual structure and constraints of the available data, thereby avoiding assumption-driven analysis that is based solely on an abstract task description.

**Fig. 1.**
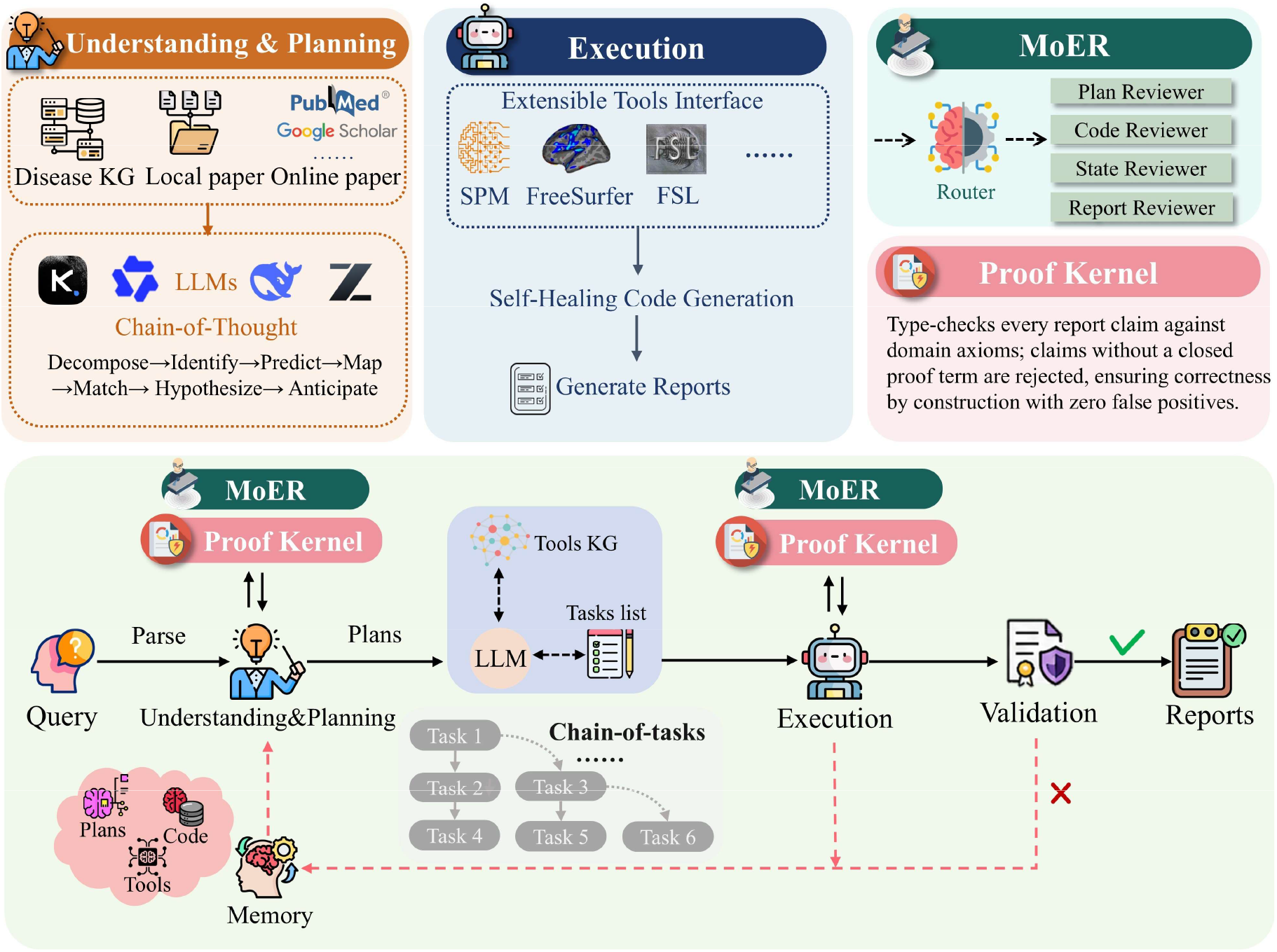
Overview of NEURA. The workflow proceeds through three phases: **(1) Understanding and planning**: NEURA integrates local and online evidence with a dynamic DiseaseKG and a ToolKG to generate biologically informed analysis plans and initialize proof skeletons. **(2) Execution**: NEURA maps plans to ordered neuroimaging toolchains, generates context-aware code and binds tool outputs to proof certificates. **(3) Validation and iteration**: MoER and the proof kernel operate across the workflow: MoER provides expert-guided methodological review, whereas the proof kernel performs deterministic verification against domain axioms and tool-derived evidence. Failed review or proof checks trigger memory-augmented reflection and workflow revision before final reporting.

In the understanding and planning phase, NEURA aims to transform a research request into a tractable and scientifically defensible analysis strategy. To achieve this goal, it combines a dual-layer retrieval-augmented generation (RAG)^30^ system with a disease-brain region knowledge graph (KG)^31^, which enables the NEURA to assemble domain-specific evidence, connects the question to establish neurobiological knowledge, and explains these resources under the PICO^32^ framework. The purpose of this phase is not only to retrieve background information but also to construct an analysis plan that is both biologically informed and methodologically appropriate. Concurrently, the proof kernel constructs a proof skeleton with typed holes at positions where tool outputs provide evidence, verifying the type-safety of the planned tool composition before execution begins.

During the execution phase, the system queries a specialized ToolKG to translate the high-level plan into a precise and ordered sequence of computational tasks. NEURA dynamically synthesizes context-specific code to execute these tasks. As each tool completes, its output is bound to a proof certificate that fills the corresponding obligation in the proof skeleton, progressively building a formal evidence chain from planning through execution. The purpose of this phase is to operationalize scientific intent into reproducible computations while preserving flexibility across heterogeneous datasets, software ecosystems and study designs. To support continual capability growth in a rapidly evolving methodological landscape, the system incorporates an extensible tool interface through which new analytical skills can be acquired directly from the literature.

In the verification phase, NEURA subjects both intermediate decisions and final outputs to methodological and factual checks through two complementary layers. The MoER module routes intermediate artefacts to domain-specific reviewers, which assess them against expert-curated normative specifications. This layer provides flexible guidance on parameter validity, statistical conventions and methodological quality. In parallel, a proof kernel acts as a deterministic gate for reportable claims: each claim is type-checked against tool-derived evidence and a bank of domain axioms, and claims without a closed proof term are excluded from the report. This design separates probabilistic methodological review from formal claim verification, avoiding the circularity of using LLMs to judge LLM-generated outputs while preserving analytical flexibility. When MoER detects a suboptimal trajectory, NEURA invokes a memory-augmented reflection loop to revise the plan, parameters or execution strategy. The underlying knowledge graphs are also updated through execution feedback, allowing the system to incorporate new evidence, tool behaviour and task outcomes as they accumulate.

Ultimately, NEURA preserves the full analytical provenance of the study, including retrieved evidence, plans, scripts, tool outputs and execution logs, and compiles these materials into a final report accompanied by a coverage triple (certified/qualified/stripped) that quantifies the formal evidentiary strength of each claim. In this way, the system is designed not only to automate neuroimaging analysis but also to render the entire process traceable, reviewable and extensible. Additional details on the system workflow are provided in the Methods section.

### 2.2 Evaluation Settings

#### Datasets

To quantitatively evaluate the performance of NEURA, we curated NeuroEval, which is a comprehensive neuroimaging tool evaluation dataset that features 110 varied analysis tasks that span multiple disease domains and complexity levels. The process of dataset construction is elaborated upon in Sec. 4.7. The final dataset was split into two levels: level 1 includes 27 tasks that can be performed with 3–4 tool steps, and level 2 comprises 83 tasks that need more than 5 tool steps with multimodal integration. This dual-level structure enables a more detailed judgement to be made about the ability of an agent to handle both simple and complex neuroimaging workflows.

#### Evaluation Metrics

To rigorously assess the effectiveness of NEURA, we used three distinct evaluation metrics: (1) planning accuracy, which is a composite metric that quantifies overall tool planning quality; (2) a tool selection score that measures the alignment between the predicted tool set and reference tools verified by human experts; and (3) a tool order score that evaluates the correctness of the tool execution sequence using longest common subsequence matching. These metrics collectively provide an assessment of the effectiveness of tool selection and the robustness of pipeline construction.

#### Baselines

To rigorously assess the efficacy of our system design, we benchmarked NEURA against a “direct LLM query” baseline. In this control setting, foundation models were supplied with identical task descriptions and tool specifications without the comprehensive agentic architecture proposed in our framework. This comparative design effectively isolates the performance gains that are attributable to our holistic system from the intrinsic capabilities of the underlying language models.

#### Foundation Models

In our experiments, we used different foundation models for the Planner, Executor and Reviewers, including the state-of-the-art large-scale models Qwen3-235B, DeepSeek-V3.2, GLM-4.6, and Kimi-K2. All the models were accessed through official APIs with standard settings. We also considered smaller but more efficient models such as Qwen2.5-7B for situations in which inference needs to occur more quickly. Because they have fewer parameters, these lightweight models have much lower latency, which makes them good for environments where real-time interactive applications are needed. We also used specialized coding models such as GLM-4.6 and the Qwen3-coder for dynamic code generation to produce executable analysis scripts. NEURA can balance planning accuracy, inference speed, and code generation quality because of this multimodal architecture. This balance ensures that our framework performs well in a variety of deployment scenarios.

#### Analysis of the Results

A performance comparison between NEURA and a direct LLM baseline across different foundation models is shown in Fig. 2a–c. Across all the tested foundation models, NEURA consistently outperforms direct LLM in terms of planning accuracy, tool selection and tool ordering. The average gains of 30.5%, 25.6% and 36.7%, respectively, indicate that the improvement was not simply due to the underlying language model but to the agentic architecture that grounds reasoning in disease knowledge, tool dependencies and validation rules. These results support the central premise of NEURA: Reliable neuroimaging workflows require structured interactions between scientific reasoning and executable tool constraints rather than ungrounded language-model planning alone.

**Fig. 2.**
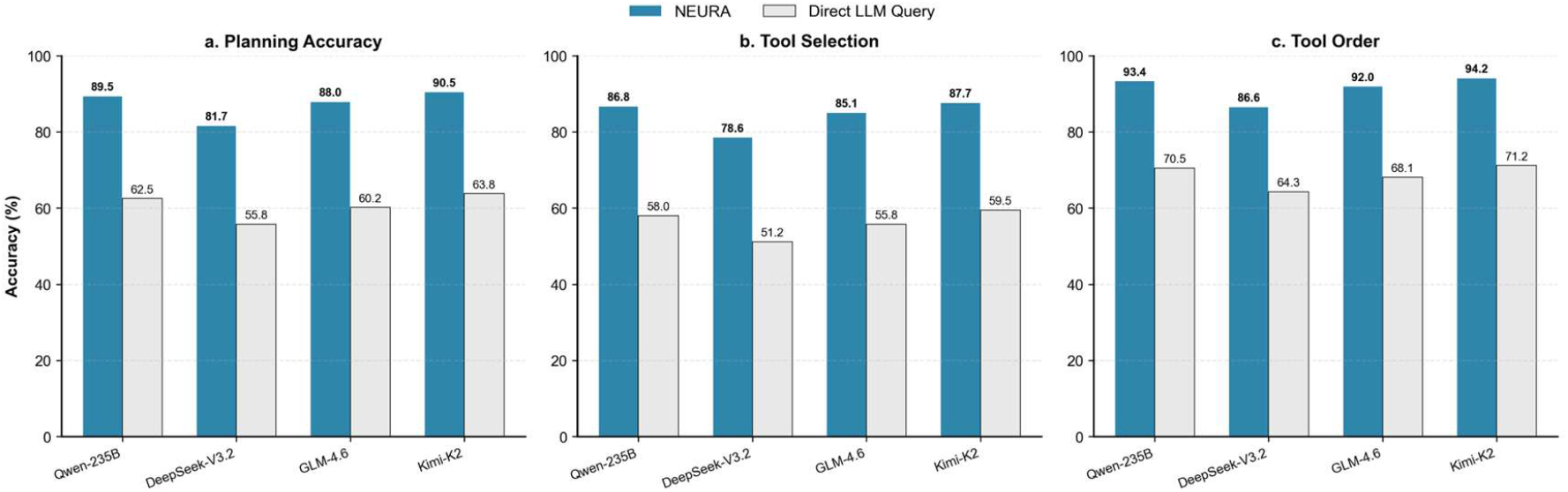
Comparison of NEURA and direct LLM queries. (a)–(c) Evaluation metrics that compare NEURA against a direct LLM query baseline across four foundation models: Qwen3-235B, DeepSeek-V3.2, GLM-4.6, and Kimi-K2. NEURA consistently outperformed the direct query approach across all the metrics, thus demonstrating the substantial efficacy of the agent architecture in orchestrating complex neuroimaging workflows compared with unassisted foundation models.

The performance of the NEURA stratified by tool category and disease type is shown in Fig. 3a–c. NEURA demonstrated consistent performance across tool categories, with most models achieving above 85% accuracy in each domain. NEURA also maintained high performance across diverse neuropsychiatric conditions. These results validate the utility of our framework for accelerating neuroimaging research workflows across diverse analysis scenarios and clinical domains.

**Fig. 3.**
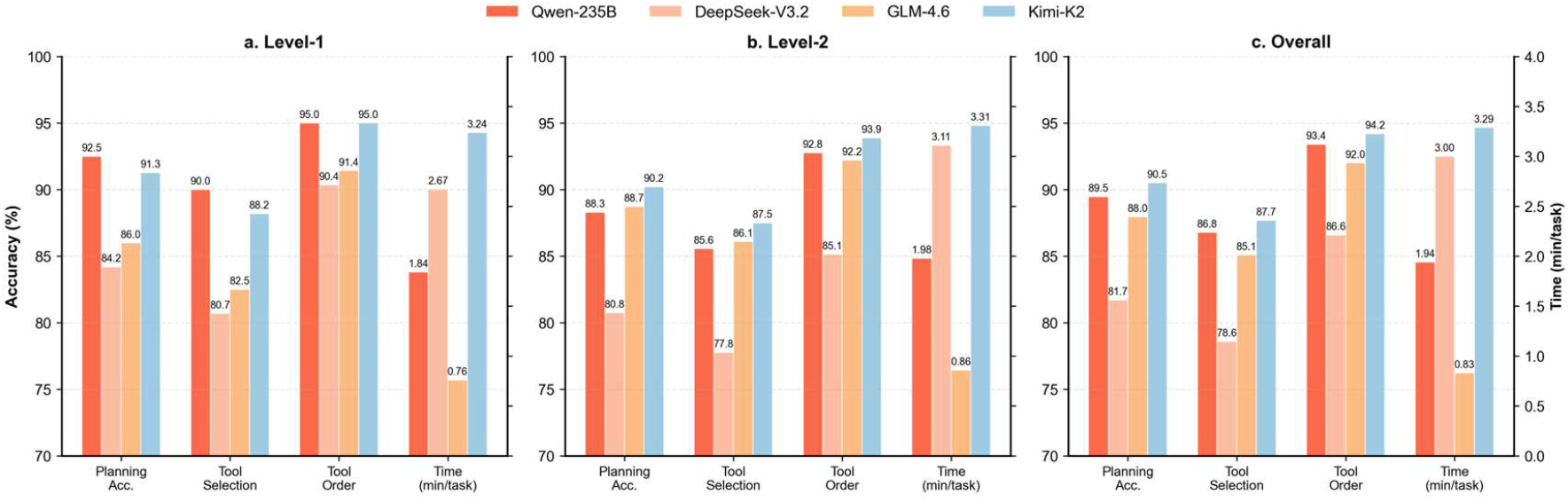
Comparison of various foundation models within NEURA. (a)–(c) Performance assessment of NEURA when powered by different state-of-the-art LLMs. (d) Comparison of the average analysis time required to complete the planning process for each model. This comparative analysis highlights the trade-offs between reasoning precision and computational latency and illustrates the adaptability of the system to different underlying architectures.

### 2.3 Evaluating proof-carrying verification with controlled hallucination injection

LLMs may generate neuroimaging reports that are fluent and internally coherent yet inconsistent with the evidential and methodological constraints of the analysis. To test whether proof-carrying verification can detect scientifically consequential errors, we designed a controlled hallucination-injection experiment. For each of the 110 NeuroEval tasks, we curated a proof-carrying report comprising three linked components: a structured report with statistical, anatomical and methodological claims; a valid proof state encoding the supporting evidence chain; and a trusted artefact registry linking tool-output hashes to their postconditions. This representation mirrors a completed NEURA execution. We introduced nine types of deterministic error(Table S1), namely, numerical falsification, anatomical substitution, methodological omission, provenance fabrication, cross-modality misapplication, statistical misuse and pipeline-order violation. Each perturbation altered the report, the proof state, or their correspondence, thereby creating ground-truth violations that require claim–evidence cross-validation. The resulting benchmark contained 990 corrupted reports and 110 clean baselines.

We evaluated two detectors: (1) the LeanNI proof kernel (zero LLM cost) and (2) an LLM-as-judge baseline (DeepSeek-V3.2 with a structured prompt). The proof kernel detected all injected errors across the nine categories, achieving precision, recall and F1 scores of 1.0 with no false positives. In contrast, the LLM judge showed high precision but limited sensitivity, detecting 50.6% of the injected errors (precision = 0.998, F1 = 0.672, false-positive rate = 0.009). Per-type analysis revealed a characteristic failure mode (Table S2): the LLM judge identified semantically obvious violations such as modality mismatch, sample-size manipulation and fabricated citations but largely missed errors that required formal comparison with the evidence state, including omitted quality-control steps, inappropriate statistical tests and falsified p-values. These results indicate that textual plausibility is insufficient for auditing scientific agents when the critical error lies in the relation between a claim and its computational provenance.

A cumulative ablation (Table S3) revealed that the kernel-type checker alone accounted for 44.4% of the injection types through postcondition verification. Adding domain-specific axiom layers (consistency, QC, modality, pipeline, and citation) increases the recall to 100%, with the false-positive rate remaining zero throughout—a property guaranteed by the axiom-grounded design.

### 2.4 Case Studies

To evaluate the effectiveness of NEURA, we applied it in clinical neuroimaging research tasks using data from patients with spinocerebellar ataxia type 3 (SCA3). SCA3 is the most common dominantly inherited spinocerebellar ataxia worldwide. It is characterized by progressive degeneration of the cerebellum, along with motor and cognitive impairments^33^. Although motor manifestations such as gait ataxia, dysarthria, and oculomotor abnormalities are hallmark features, neuroimaging studies have consistently demonstrated widespread structural alterations, including cerebellar atrophy, brainstem degeneration, and subcortical grey matter loss^34,35^.

#### Data Acquisition

We recruited 44 patients with genetically confirmed SCA3 and 60 age-matched healthy controls (HCs) from the neurology outpatient clinic. All MRI data were obtained on the same day as the clinical evaluation. MRI scans were acquired on a Siemens (Erlangen, Germany) 3T Trio TIM MRI system equipped with a standard Siemens 8-channel head coil. All the subjects were instructed to keep their eyes closed, relax, think of nothing in particular, and avoid falling asleep (as confirmed by all the participants immediately after the scan). The protocol included three-dimensional T1-weighted anatomical, diffusion and resting-state functional MRI. T1-weighted structural imaging was performed using a 3D magnetization-prepared rapid-acquisition gradient echo sequence with the following parameters: repetition time = 1900 ms, echo time = 2.52 ms, flip angle = 9°, slice thickness = 1 mm, field of view = 256 mm × 256 mm, matrix = 256 × 256, and voxel size = 1 mm × 1 mm × 1 mm, with 176 slices and no gap. Diffusion imaging acquisition was performed using an echo planar imaging (EPI) sequence with the following parameters: repetition time = 8000 ms, echo time = 80.8 ms, matrix = 128 × 128, voxel size = 2 × 2 × 2 mm, slice thickness = 2 mm, no slice gap, b = 0 and 1000 s/mm^2^, 64 directions and a total of 67 slices. Additionally, all the participants completed the Montreal Cognitive Assessment (MoCA) to evaluate global cognitive function^36^.

#### 2.4.1 NEURA Facilitates Structural Brain Analysis

Voxel-based morphometry (VBM)^37^ analysis is a basic approach for investigating differences in the volume of grey matter in different areas between groups of patients and HCs. In traditional VBM workflows, researchers have to manually coordinate several preprocessing steps, decide in real time which tools and parameters to use on the basis of intermediate results, and refine analyses repeatedly when initial attempts show problems. NEURA automates this intricate iterative procedure using autonomous planning, execution, reflection, and replanning cycles.

Given a user query (Fig. 4), NEURA initiated an autonomous structural MRI workflow that ran for 27 hours across two planning–execution–validation cycles. The system combined voxelwise and surface-based analyses, using SPM with DARTEL^38^ for high-dimensional normalization, the spatially unbiased infratentorial template (SUIT) atlas^39^ for cerebellar parcellation and FreeSurfer for complementary cortical and subcortical morphometry. NEURA selected statistical tests on the basis of data distribution checks, enforced Benjamini–Hochberg FDR correction^40^ and detected missing MoCA scores, after which it adjusted the sampling strategy to maintain complete-case consistency.

**Fig. 4.**
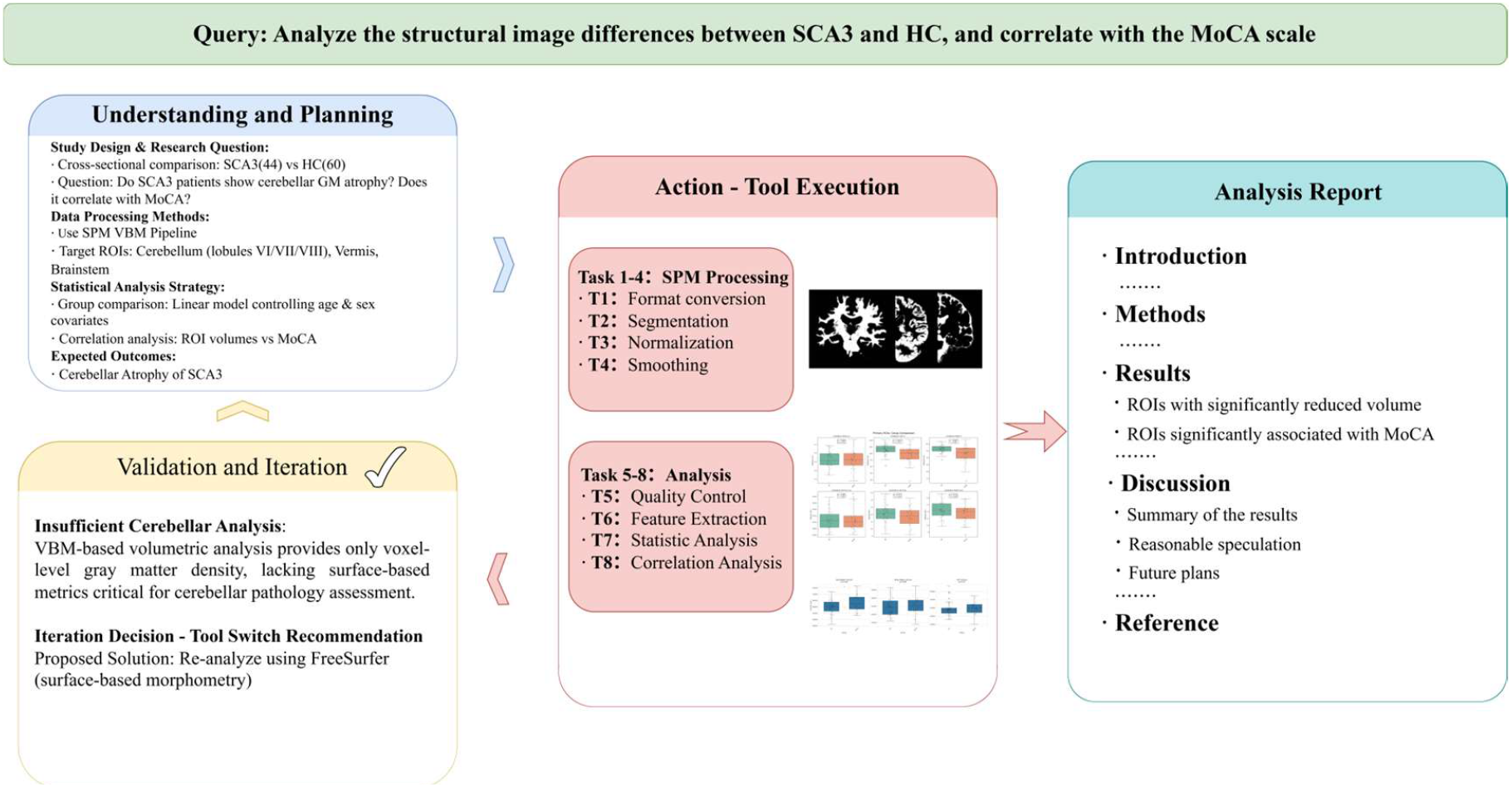
Workflow and results of structural image analysis using NEURA

After two complete planning–execution–validation cycles, NEURA converged onto stable, validated results (Table S4 and Fig. 4). The automated analysis correctly revealed brain atrophy patterns consistent with those of established SCA3 pathology, with widespread grey matter volume reductions across cerebellar subregions. The most pronounced differences were observed in bilateral cerebellar lobules V and VI, with the left lobule VI exhibiting the greatest effect size (Cohen’s *d* = 0.73), accompanied by significant atrophy in the somatomotor-related anterior lobes (I–IV). With respect to clinical correlations, NEURA revealed a positive association between left Crus I volume and MoCA scores (*r* = 0.30). These findings align with known neuropathology in which cerebellar cortical degeneration particularly affects lobules V–VI^41^. VBM meta-analyses have consistently revealed significant volume reductions in cerebellar hemispheric lobules III–VI in SCA3 patients^34^, with lobule VI volume loss specifically associated with cognitive impairment^41^.

To corroborate the automated findings, three independent neuroimaging experts performed parallel manual analyses using standard processing pipelines. This human-derived benchmark substantiated the accuracy of the NEURA results. Critically, the system executed the complete workflow with full autonomy. It successfully navigated technical impediments such as incomplete preprocessing artefacts, missing clinical covariates, and the requirement for adaptive statistical test selection without any human intervention.

#### 2.4.2 NEURA facilitates DTI analysis

Diffusion tensor imaging (DTI) provides quantitative measurements of white matter microstructure through the characterization of water diffusion properties within neural tissue^42^. Key DTI metrics include fractional anisotropy (FA), which reflects the directionality of diffusion and is sensitive to axonal integrity, and mean diffusivity (MD), axial diffusivity (AD), and radial diffusivity (RD), which characterize the magnitude and orientation of water diffusion. Previous studies have reported white matter abnormalities in the SCA3, particularly in the brainstem and cerebellar peduncles^43^. To demonstrate the ability of NEURA to perform diffusion imaging analysis, we applied it to DTI data from the same cohort. Owing to missing modality acquisitions, the sample size was reduced to 49 HCs and 42 SCA3 patients.

Given a user query (Fig. 5), NEURA first parsed the research question and formulated an analysis plan that included preprocessing, tensor fitting, spatial normalization, and statistical comparison. NEURA then orchestrated a multitool pipeline to generate voxelwise FA, MD, AD, and RD maps. For the voxelwise analysis, NEURA implemented tract-based spatial statistics (TBSS) ^44^ to project FA data onto a mean white matter skeleton, and QSDR reconstruction from DSI Studio was used for whole-brain metric extraction in MNI space. Critically, an autonomous quality control module monitored each stage for excessive head motion and processing failures. During execution, NEURA identified one outlier with aberrant FA values and autonomously excluded it; thus, the final sample was adjusted to 90 subjects (49 HCs and 41 SCA3 patients). The complete pipeline required coordination across the FSL and DSI Studio platforms, which demonstrated the ability of NEURA to seamlessly integrate multiple tools.

**Fig. 5.**
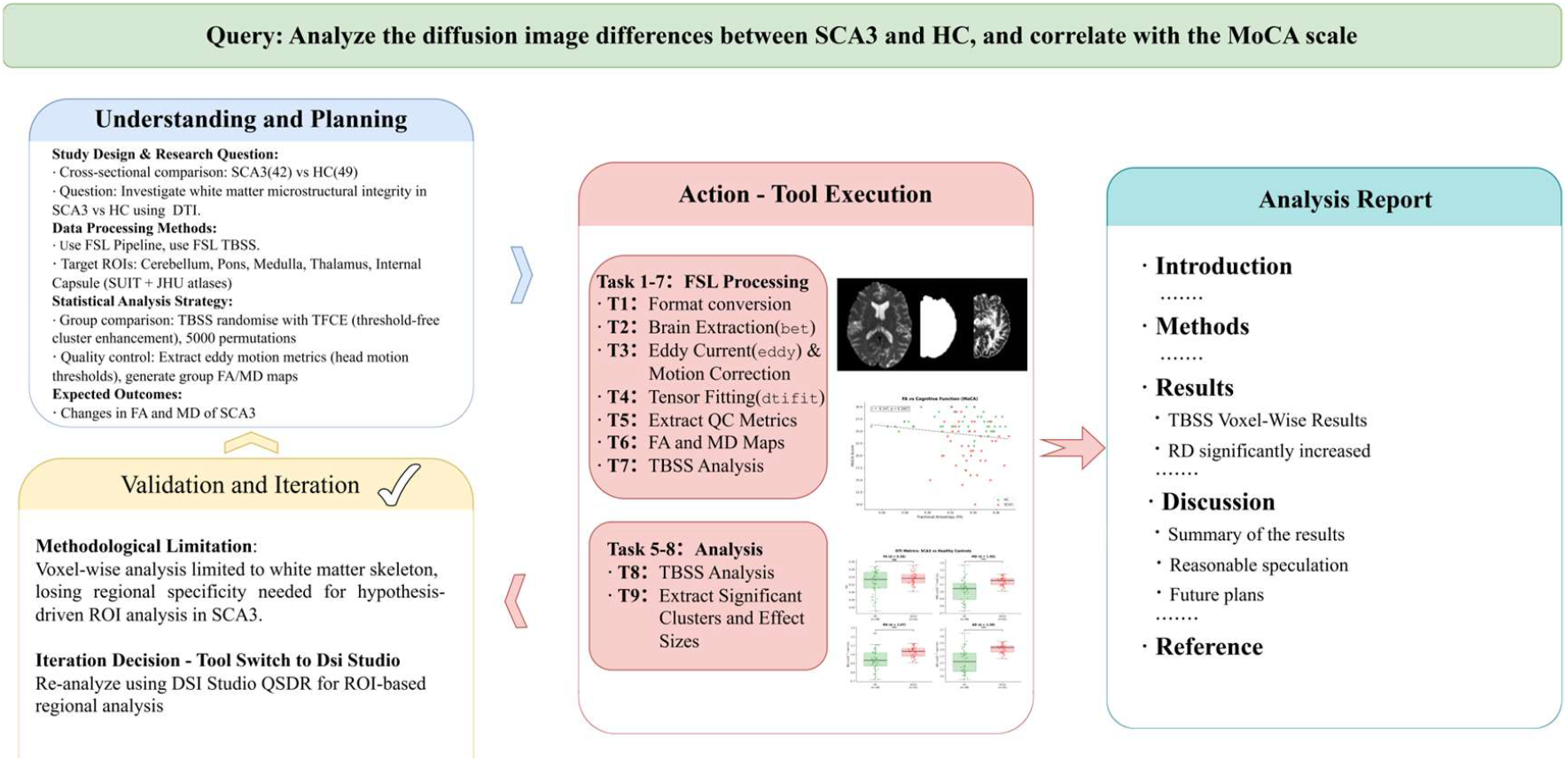
Workflow and results of DTI analysis using NEURA

Group comparisons revealed significantly elevated diffusivity metrics in SCA3 patients (Table S5). MD, RD, and AD all showed large effect sizes (Cohen’s d > 1.0; all p < 0.0001), whereas whole-brain FA did not differ significantly between groups (p = 0.079), which likely reflected global averaging across regions with heterogeneous pathology. These findings complement the grey matter atrophy observed in Case 1 and align with previous DTI studies that reported increased diffusivity in SCA3. Similarly, the analysis results were manually processed and analysed by three neuroimaging experts using traditional methods to verify their accuracy.

In summary, these two case studies demonstrate the ability of NEURA to perform complex neuroimaging analyses through autonomous planning, multitool collaboration, iterative improvement, and rigorous result validation.

## 3. Discussion

NEURA reframes the role of agentic AI in neuroimaging. Rather than treating AI as an image classifier, a code generator or a pipeline executor, NEURA acts as a governed research agent that connects biological hypotheses, computational execution and methodological accountability. This distinction is important because the central challenge in modern neuroimaging is not only in larger, multimodal and more heterogeneous datasets^1,45^, but also in converting increasingly large and heterogeneous imaging resources into evidence that is interpretable and reviewable.

Existing workflow standards and engines such as Nipype^14^ and BIDS Apps^46,47^ have been essential for standardizing data organization, preprocessing environments and predefined neuroimaging pipelines. However, they generally require users to specify the analysis strategy in advance. NEURA addresses a different level of the research process. It accepts natural-language scientific questions, inspects the available workspace and constructs analysis strategies under biological, computational and statistical constraints. Compared with general-purpose scientific agents such as Coscientist^25^ and ChemCrow^25^ and biomedical agentic systems such as DeepRare^17^, NEURA demonstrates how domain adaptation and formal verification can be combined in a scientific agent. DiseaseKG, ToolKG and literature-derived skills provide the structure needed to translate neuroimaging questions into executable workflows. The proof kernel serves a distinct function: it checks whether each reportable claim is supported by explicit computational evidence under the specified axiom bank and trust assumptions.

The evaluations support both aspects of this design. On NeuroEval, NEURA achieved a planning accuracy of 89.5% and outperformed direct LLM queries by 30.5% in planning accuracy, 25.6% in tool selection and 36.7% in tool ordering, indicating that domain grounding improves workflow construction. The hallucination-injection experiment further reveals why verification cannot be delegated to an LLM judge alone. DeepSeek-V3.2 detected only 50.6% of injected errors (F1 = 0.672), performing well on semantically obvious violations but missing errors that required comparison with the underlying evidence, including omitted quality-control steps, inappropriate statistical tests and falsified p-values. In contrast, the proof kernel detected all injected error classes without false positives by testing claim–evidence consistency rather than textual plausibility. Finally, the case studies show that this architecture can support end-to-end structural and diffusion MRI analyses, producing findings consistent with those of established pathology and independent expert analyses.

Several limitations remain. First, NEURA’s guarantees are bounded by the coverage and quality of its curated knowledge resources. The current proof kernel uses 136 expert-defined axioms covering 8 neurological conditions and 6 major analysis tools, and its evolving knowledge graphs depend on reliable literature retrieval, evidence extraction and validation. Claims outside this coverage are qualified or stripped rather than certified, making expert-reviewed expansion of the axiom bank essential for broader use. Second, the kernel verifies consistency between claims, axioms and trusted tool outputs, but it does not prove the internal correctness of external software or the validity of the acquired data. Erroneous tool outputs or uncaptured data-quality problems may therefore propagate into certified claims, underscoring the need for stronger tool validation, richer quality-control axioms and expert oversight in high-stakes settings. Third, larger external evaluations are needed. NeuroEval measures planning quality across literature-derived tasks, and the SCA3 studies show end-to-end feasibility, but validation across population-scale cohorts, sites, modalities and disease spectra remains necessary. Future work should expand and version the axiom bank, validate reasoning chains with experts, perform module-wise ablation and failure-mode analyses, and develop stronger formal back ends for machine-checked verification.

Beyond neuroimaging, NEURA illustrates a general strategy for building trustworthy scientific agents: separate the generation of workflows and claims from their verification. The planning and execution components can remain flexible and LLM-driven, whereas reportable claims must be grounded in explicit evidence and checked against a domain-specific axiom bank. This architecture could be adapted to other data-intensive fields by replacing the neuroimaging axioms with field-specific disease or entity knowledge, tool contracts, statistical assumptions and reporting standards. Thus, although NEURA is instantiated here for neuroimaging, its proof-carrying design provides a template for scientific agents in which flexibility and auditability are not treated as opposing goals.

Although NEURA accelerates research, it functions as an assistant rather than a replacement for human scientists. By automating complex neuroimaging processing and analysis workflows while rigorously maintaining human-in-the-loop oversight, the system empowers researchers to shift from technical implementation to higher-level hypothesis generation. We strongly advocate for transparent disclosure of such AI assistance^49^ to ensure that this paradigm shift accelerates scientific discovery without compromising the integrity that remains the hallmark of human research.

## 4. Methods

In this section, the methodology of NEURA, including the implementation details for the overall workflow and the associated LLMs, is presented.

### 4.1 Problem Formulation

We formalize the neuroimaging research automation task as a constrained optimization problem. Given a natural language research question *Q* and a neuroimaging dataset 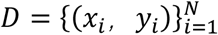, where *x*_*i*_ represents multimodal brain images and *y*_*i*_ denotes subject metadata, the system produces three outputs: a scientific report ℛ in IMRAD format, executable analysis scripts 𝒮, and validated statistical results 𝒱.

The optimization objective maximizes scientific quality while satisfying a hallucination constraint:

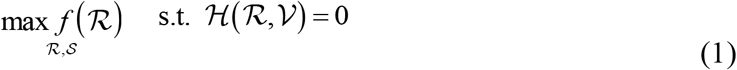

where *f* (ℛ) quantifies the report quality through structure completeness, statistical rigor, and logical coherence and ℋ (ℛ,𝒱) represents the hallucination constraint, which ensures that every claim in ℛ is grounded in the computational results 𝒱.

### 4.2 Main Workflow

The workflow is structured into three distinct phases: semantic understanding and planning, compositional execution, and validation and iteration. Drawing on principles from cognitive architectures for language agents^24^, this tripartite framework systematically transforms natural language queries and raw data into evidence-grounded, executable analyses.

During the **semantic understanding and planning phase**, the system parses the natural language query from the user into a structured analysis specification and extracts critical variables, including target phenotypes, imaging modalities, study cohorts, and comparative conditions. By leveraging a dual-layer retrieval mechanism and a dynamic DiseaseKG, NEURA synthesizes supporting evidence to propose candidate brain regions and effectively grounds disease descriptors in pathology-informed neuroanatomical associations. This specification is subsequently translated into an actionable research plan that adheres to the PICO ^32^ framework and is operationalized as an ordered tool sequence. For each stage, the system configures modality-compatible workflows with precise parameters and defined intermediate artefacts (e.g., preprocessed images and statistical maps), thereby yielding an executable plan that has been rigorously validated against input constraints.

During the **compositional execution phase**, the system transforms the research plan into a precise workflow via a specialized ToolKG. This graph maps the planned steps to specific software dependencies and ensures input/output compatibility. To accommodate novel methodologies, NEURA uses an extensible “skill” learning interface that enables it to autonomously distill reusable analytic parameters directly from the methodological literature. NEURA dynamically synthesizes context-specific code to orchestrate these heterogeneous tools and creates a valid execution chain that persists all intermediate artefacts and logs to a shared workflow state.

To guarantee the reliability of the aforementioned processes, a MoER mechanism operates concurrently across the planning and execution phases. This module routes intermediate outputs to domain-specific experts. By enforcing strict evaluation criteria at every decision point, the system ensures methodological rigor and syntactic correctness prior to final validation.

During the **reflective iteration phase**, if validation fails or if the overall quality score falls below the acceptance threshold, the system triggers reflection, revises the plan or parameters, and repeats execution until the criteria are met or a maximum number of iterations is reached.

These phases are implemented by cross-cutting components for structured reasoning, reflection-driven refinement, and evidence-based auditing, which are described in the subsequent sections.

### 4.3 Semantic Understanding and Planning

In this section, the functional modules that operationalize the semantic understanding and planning phase, spanning structured question interpretation, evidence retrieval, and skill reuse, are described.

#### 4.3.1 Structured Reasoning Protocol

To enable explicit, stepwise reasoning from an input research question to a structured analysis specification, the reasoning agent applies a seven-step chain-of-through protocol^22^. The protocol comprises the following: (1) problem decomposition: decomposing the research question into subquestions; (2) disease identification: extracting the mentioned or implied neurological conditions; (3) brain region prediction: suggesting affected regions on the basis of pathology-informed associations; (4) concept mapping: translating clinical terminology into neuroimaging constructs; (5) method matching: identifying appropriate analyses; (6) hypothesis formation: generating testable predictions; and (7) issue anticipation: identifying methodological challenges and constraints that may affect downstream execution. Crucially, to balance exploratory reasoning with strict adherence to the protocol, this process is orchestrated via a multimodel allocation strategy, where higher-temperature models handle initial semantic parsing and lower-temperature models enforce logical consistency in the final specification.

#### 4.3.2 Knowledge Infrastructure

In this section, the knowledge infrastructure that enables evidence-based research automation, from the literature retrieval to reusable skill accumulation, is described.

##### (1) Dual-Layer Retrieval-Augmented Generation

To provide both stable domain knowledge and access to up-to-date external evidence for task-specific reasoning, NEURA implements a dual-layer RAG^30^ framework that combines local knowledge bases with real-time literature search. For the local layer, documents are embedded into a shared semantic space:

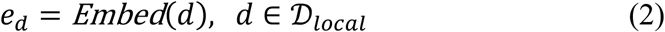

Given a query q, Retrieval computes the cosine similarity and returns the top-k documents:

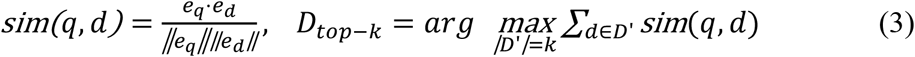

The real-time layer retrieves additional evidence by querying PubMed and extracting titles, abstracts, authors, journals, publication years and DOIs. Local chunks and PubMed records are merged to guide downstream neuroimaging workflow selection and parameterization.

##### (2) Dynamic Disease-Region Knowledge Graph

To ground neuroimaging reasoning in structured neuropathological knowledge and support biologically informed analytical planning, a key innovation of NEURA is the disease–brain KG designed by eight neuroimaging experts, which encodes pathological region associations. In the knowledge graph 𝒢_disease_ = (*V*_disease_, *V*_region_, *E*_assoc_, *W*), the nodes represent diseases and brain regions, the edges represent associations, and the weights *W* capture association strength and are derived from evidence from the literature.

Initially, the graph encodes established mappings for eight neurological conditions. Spinocerebellar ataxia involves the cerebellum, brainstem, and basal ganglia. Similar mappings exist for Alzheimer’s disease, Parkinson’s disease, amyotrophic lateral sclerosis, major depressive disorder, schizophrenia, ADHD, and autism spectrum disorder. Critically, this knowledge graph supports dynamic updates during conversational interactions, which enables it to incorporate information derived from system usage and newly accessed literature. When the system processes new research questions or retrieves novel findings from the literature, the graph evolves through three mechanisms:

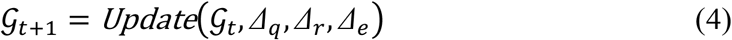

where *Δ*_*q*_ represents query-driven updates, *Δ*_*r*_ denotes retrieval-driven updates (relationships extracted from retrieved literature), and *Δ*_*e*_ denotes execution-driven updates (statistically significant findings from completed analyses).

The associated weights can be updated as follows:

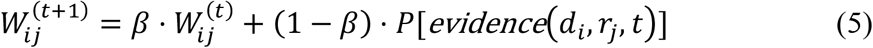

where *P*[⋅] ∈ [0,1] denotes the confidence score of the evidence and *β* = 0.95 balances prior knowledge with new evidence. This update enables the system to accumulate domain knowledge across sessions, thereby progressively improving region recommendations as more research tasks are processed.

##### (3) Literature-Derived Skill Library

To support methodological extensibility and continual adaptation to the evolving neuroimaging literature, the system automatically extracts reusable analysis skills from the neuroimaging literature. An analysis skill *S* comprises four components:

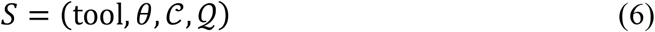

where tool represents the analysis software,θ denotes the parameter configuration, 𝒞 represents the applicability context (disease and modality), and 𝒬 captures the quality indicators from source publications.

The extraction pipeline implements the processes described in the Methods section in five stages: task identification using structural cues, tool recognition via named entity recognition, parameter extraction by parsing numerical values and thresholds, context encoding to capture disease and modality, and quality assessment on the basis of effect sizes and validation metrics.

##### (4) Semantic Skill Matching

When a research task arrives, semantic matching recommends appropriate skills. Both tasks and skills are embedded into the shared space:

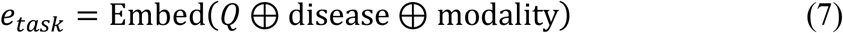

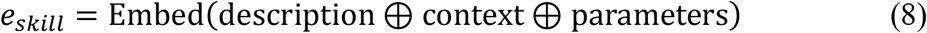

where ⊕denotes concatenation.

The matching score integrates semantic similarity with constraint satisfaction:

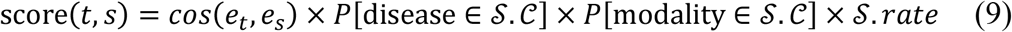

where *P*[⋅] ∈ [0,1] quantifies the applicability confidence on the basis of the semantic similarity between the task context and the validated domains of the skill and 𝒮. *rate* reflects historical success.

##### (5) Context-Aware Skill Adaptation

When learned skills are applied to new contexts, parameter adaptation addresses source–target differences:

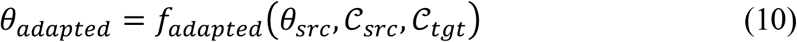

where the adaptation function *f*_*adapted*_ accepts three inputs to produce context-appropriate parameters, θ_*src*_ denotes the original parameters extracted from the literature, 𝒞_*src*_ denotes the source context where the skill was validated, and 𝒞_*tgt*_ denotes the target context for the current research task.

Postexecution validation updates the skill quality via an exponential moving average:

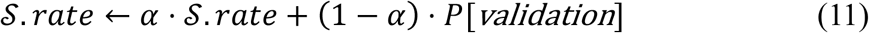

where *P*[validation] ∈ [0,1] reflects the validation quality, with α enabling continuous learning from accumulated experience.

### 4.4 Compositional Execution

Herein, we explain how our system selects the right tools, builds analysis pipelines, and runs them across different platforms.

#### 4.4.1 Tool Knowledge Graph

To translate high-level analytical plans into valid, executable tool sequences, we model neuroimaging tools as a knowledge graph 𝒢_*tool*_ = (*V, E*_*dep*_, *E*_*io*_, *E*_*cat*_), where *V* represents tools, *E*_*dep*_ captures dependencies, *E*_*io*_ represents input/output compatibility, and *E*_*cat*_ encodes categorical relationships.

The schema encodes five information categories: functional descriptions, format specifications, categorical labels (preprocessing, analysis, visualization, and QC), software provenance, and safety annotations for irreversible operations. Three dependency types are formalized. Sequential dependencies occur when the output of tool *T*_*i*_ serves as an input for *T*_*i+1*_:

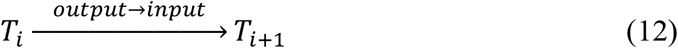

Parallel execution enables tools to run simultaneously on independent streams:

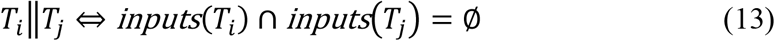

Conditional execution is contingent on upstream evaluation:

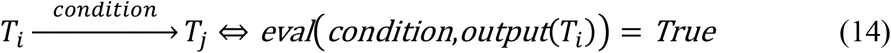

Similarly, the tool knowledge graph is designed as an open and extensible architecture and is not limited to the currently implemented toolset. Tool extensibility is achieved through two complementary mechanisms. First, literature-driven automatic discovery is performed: When new publications are being processed, the system applies named entity recognition (NER) to identify candidate tools not yet in the graph:

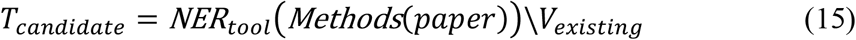

The extracted tools are enriched with parameter configurations from the same source and queued for validation before graph integration.

Second, user-defined tool registration is conducted: Researchers can register custom tools by providing interface specification files. After registration, the system automatically infers connectivity by matching input/output formats:

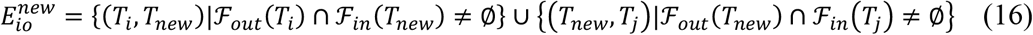

where 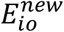 denotes the set of new input/output edges that connect the newly registered tool to existing tools, ℱ_*out*_(*T*_*i*_) denotes the set of output formats produced by tool *T*_*i*_, and ℱ_*in*_(*T*_*new*_) denotes the set of input formats accepted by tool *T*_*new*_. This automatic edge inference ensures that newly registered tools are immediately available for pipeline construction without manual dependency specification. The system checks whether the output of each existing tool matches the input requirements of the new tool and automatically establishes connectivity when compatibility is detected.

A unified abstraction layer supports both the Windows and WSL2 Ubuntu environments. The system automatically detects the locations of tool installations and route requests.

#### 4.4.2 Graph-Guided Tool Selection

For tool selection, four-step graph-based retrieval is used. First, by full-graph retrieval, semantically similar tools are identified:

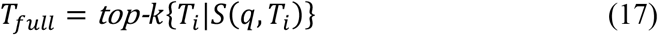

where *S* (*q, T*_*i*_) computes the embedding similarity between the query and tool descriptions. Second, subgraph exploration is conducted to identify related tools:

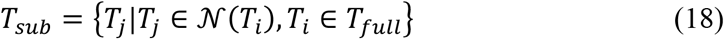

where 𝒩 (*T*_*i*_) denotes the neighbourhood set that contains all the tools directly connected to tool *T*_*i*_ in the knowledge graph. Intuitively, this step finds all tools that are adjacent to the initially selected tools, thereby capturing functionally complementary tools. For example, if FSL registration (FLIRT) is selected, the neighbourhood includes downstream tools such as FEAT and preprocessing tools. Third, combination and reranking produces the final candidate set:

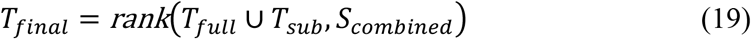

Fourth, chain-of-tools generation is conducted to synthesize the optimal sequence with respect to dependencies.

#### 4.4.3 Self-Healing Code Generation

Diverging from template-based paradigms, the execution framework relies on dynamic, context-aware code synthesis. The system generates bespoke scripts tailored to the specific analytic constraints of each task and embeds them within a self-correcting closed-loop protocol. This mechanism enforces robustness through iterative refinement: generated code is first validated by abstract syntax tree (AST) parsing and subsequently subjected to execution monitoring. Upon failure, the system triggers an error-informed regeneration cycle and systematically traverses a hierarchical error taxonomy that resolves syntactic violations, dependency conflicts, runtime exceptions, and data compatibility mismatches.

Furthermore, to optimize the computational throughput, we implement a content-addressable memoization layer. By cryptographically hashing task descriptors and input states, the system retrieves cached metadata for recurrent operations, thereby eliminating redundant computations.

#### 4.4.4 Proof-Carrying Verification

To provide deterministic anti-hallucination guarantees without LLM self-evaluation, we designed a formal verification layer inspired by the Curry-Howard correspondence and proof-carrying code. Every claim in a neuroimaging report, whether a statistical finding, a methodological assertion, or a tool-output reference, is modelled as a proposition; the supporting evidence (tool execution records, axiom instantiations, inference steps) is modelled as a proof term. A claim is accepted if and only if its proof term type-checks in a trusted kernel.

##### (1) Proof Kernel Design

The trusted core implements a single recursive function that type-checks a proof term against an expected proposition. Four proof-term constructors are recognized:

- **AxiomRef:** References a registered axiom with variable instantiation. The kernel verifies registry membership, recursive premise checking, and the absence of unbound variables.
- **ToolOutputRef:** A tool execution certificate. The kernel verifies that the artifact’s hash appears in the trusted context, grounding the claim in an actual computation.
- **RuleApp:** Applies an inference rule to child-proof terms, for example to verify valid tool composition, confirm that a statistical threshold has been met. The kernel recursively checks all the child proof terms before performing rule-level unification.
- **Hole:** An open proof obligation. Any term containing a hole is unconditionally rejected; this is the mechanism by which unsubstantiated claims are captured.

##### (2) Domin Axiom Bank

The kernel operates over 136 curated axioms in four categories:

- **Disease–region mappings (84):** Expert-curated axioms encoding established associations between neurological conditions and brain structures. Each axiom is assigned a confidence tier and linked to supporting DOI-cited evidence, allowing the verifier to distinguish well-established disease–region relationships from weaker or heuristic associations.
- **Tool contracts (27):** Axioms extracted from tool documentation, usage specifications and expert-curated operating rules, including input–output type signatures, modality constraints, parameter requirements and pipeline dependencies. For example, these axioms specify whether a tool accepts T1-weighted, diffusion or BOLD inputs; what outputs it produces; and which preprocessing steps, such as eddy-current correction before tensor fitting, must precede downstream analysis.
- **Statistical prerequisites (15):** Expert-curated axioms that encode statistical assumptions and reporting requirements, including normality checks for parametric tests, correction for multiple comparisons, and minimum sample-size constraints.
- **Method applicability (10):** Expert-curated axioms that define which analytical methods are appropriate for specific study designs, imaging modalities and research objectives.

The axiom bank is extensible and version controlled, allowing new or revised axioms to be incorporated as evidence, tools and methodological standards evolve, subject to expert review, evidence linkage and consistency checking.

##### (3) Proof Construction Lifecycle

NEURA is used to construct proof evidence incrementally throughout the workflow. At the planning stage, it specifies the evidence required for each planned operation and checks whether the proposed tool sequence satisfies the modality, dependency and methodological constraints before computation begins. During execution, each completed tool registers its output artefacts, hashes and postconditions, thereby linking subsequent claims to their generating computations. At reporting, each claim is audited against the completed proof state and classified as certified, qualified or stripped according to the strength of its supporting evidence. Claims grounded in high-confidence axioms are certified; those relying on heuristic axioms are reported with caveats; and claims without sufficient formal support are removed from the final report. This graded verification scheme provides a calibrated confidence signal and allows transparent degradation when the axiom bank is incomplete.

##### (4) Normative Specification Layer

The proof kernel operates alongside a MoER mechanism. The MoER routes intermediate artefacts to domain-specific reviewers for planning, coding, statistics and reporting, where they are evaluated against expert-curated normative specifications. This layer is designed to improve the scientific quality of the workflow by assessing whether the planned analyses, parameter choices, statistical procedures and reporting structure are methodologically appropriate.

The proof kernel plays a distinct role. Rather than judging whether an analysis is optimal or well designed, it verifies whether each reportable claim is correct with respect to explicit evidence, domain axioms and trusted tool outputs. Claims are retained only when their supporting proof is complete; unsupported or inconsistent claims are removed before reporting. This separation allows NEURA to use expert-guided review to make analyses more reasonable and rigorous, while relying on formal verification to safeguard the correctness of factual claims.

### 4.5 Reflective Iteration

In the final phase, namely, the validation and iteration phase, the workflow applies a rigorous quality control mechanism. Whenever validation metrics indicate suboptimal performance or logical inconsistencies, the system autonomously triggers a reflective optimization loop. Central to this adaptive process is the reflection agent, which operationalizes the Gibbs reflective cycle^50^ to facilitate systematic learning from execution trajectories. The protocol unfolds through a structured six-phase cognitive sequence: First, state reconstruction (Description) and uncertainty quantification (Feelings) are performed. Next, performance adjudication (Evaluation) and root-cause diagnostics are conducted via a recursive ‘5-Why’ inference mechanism (Analysis). These insights are subsequently synthesized into actionable heuristics (Conclusion), which culminates in the generation of stratified remediation strategies (Action Plans). Crucially, this mechanism enables NEURA to transcend simple parameter tuning; depending on the diagnosis, it can dynamically refine the experimental design, reorient the research trajectory, or adopt superior analytical methods. This feedback cycle is repeated until the identified shortcomings are resolved and rigorous scientific standards are met, thereby ensuring continuous self-optimization.

### 4.6 Human𝒢Agent Collaborative Control

Unlike batch systems, NEURA supports streaming execution with real-time intervention. Each node emits state updates upon completion, thereby enabling monitoring of the dashboards without polling. User intervention is possible at five control points: after plan generation (approve, modify, and replan), before tool execution (override, adjust, and skip), during long tasks (pause, resume, cancel, and checkpoint), after tool completion (accept, retry, and modify), and before report finalization (edit, regenerate, and extend). When users intervene, a formal model ensures consistency:

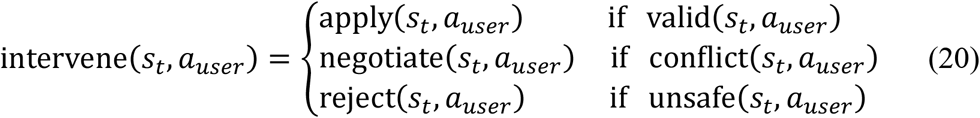

where *s*_*t*_ denotes the complete workflow state at time *t*, including the results, data, parameters, and context information of all the executed steps, and *a*_*user*_ denotes the intervention actions or requests proposed by the user at the human–computer interaction point. The system preserves the state before modifications, identifies affected downstream steps, and selectively re-executes only invalidated operations.

### 4.7 Construction of NeuroEval

To evaluate the performance of NEURA, we constructed NeuroEval, a comprehensive benchmark that comprises 110 diverse neuroimaging analysis tasks. The curation of NeuroEval followed a strict, expert-driven process that was meant to ensure that it was both scientifically sound and useful in real life.

#### (1) Curation methodology

We assembled a multidisciplinary panel of eight neuroimaging specialists to curate the dataset. Guided by a systematic extraction protocol, the experts selected high-quality publications that featured complex, multimodal workflows specific to their subfields. Methodological details were distilled from these sources into standardized task descriptions and precise research inquiries. Finally, to ensure reliability, all the cases were validated through a rigorous peer-review process.

#### (2) Dataset stratification

NeuroEval stratifies tasks into two tiers of complexity. Level 1 (27 cases) comprises intermediate workflows that involve 3–4 discrete tool operations, whereas Level 2 (83 cases) targets advanced scenarios that require more than 5 steps and multimodal integration. To evaluate clinical generalization, the benchmark encompasses 11 distinct neurological and psychiatric domains, including those related to neurodegenerative, cerebrovascular, and developmental disorders. We uniformly allocated 10 cases per category to ensure a balanced assessment across diverse pathologies.

#### (3) Standardized metadata

For each entry, a standardized schema was used to support reproducible evaluation. The metadata included (1) specific research questions, (2) a validated “ground truth” reference toolchain, (3) the source publication, and (4) specific annotations for complexity, modality, and anatomical regions. This structured format enabled precise, quantitative assessment of the ability of NEURA to translate clinical questions into executable computational pipelines.

## Data availability

The data that support the findings of this study are available on request from the corresponding author. The clinical neuroimaging and cognitive data associated with case studies are not publicly available because of genetic privacy restrictions and institutional ethical guidelines regarding patient confidentiality. NeuroEval and the formal verification resources generated for this study, including the domain axiom bank, hallucination-injection specifications and evaluation metadata, are publicly available in the NEURA GitHub repository at https://github.com/NeuroScienceLab/NEURA.

## Code availability

Code for the NEURA system and for the NeuroEval benchmark evaluations is available in our GitHub repository at https://github.com/NeuroScienceLab/NEURA.

## Acknowledgements

We gratefully acknowledge the study participants, as well as the developers of the neuroimaging tools and large language models who made this work possible. This work was supported by the Brain Science and Brain-like Intelligence Technology-National Science and Technology Major Project under grant no. G072022ZD0208903, the Natural Science Foundation of China under grant no. 62503091, and the Sichuan Province Science and Technology Foundation under grant no. 2025ZNSFSC1758.

## Author contributions

J.X., J.G. and H.C. conceptualized the study. J.X. implemented NEURA and ran it on the case studies. C.L., X.W., X.L. and Y.M. developed NeuroEval. J.W. benchmarked NEURA against the base LLMs. C.L. and J.G. evaluated the agent’s case study results. H.C. supervised the study. All the authors contributed to writing the paper.

## Ethics approval

The study was approved by the Ethics Committee of the First Affiliated Hospital of Army Medical University (reference: KY201863, KY2020191, and KY2023046). All participants provided written informed consent in accordance with the Declaration of Helsinki.

