## Supplemental Data 1 for "NEURA: A proof-carrying framework for hallucination-resistant neuroimaging automation"

### Supplementary Information

Table S1: Hallucination Injection Types

| | Injection Type | Description | Example | Detection Layer |
|---|---|---|---|---|
| 1 | swap_p_value | Falsify p-value | p=0.0000001 → p=0.12 | Kernel postcondition |
| 2 | swap_n | Falsify sample size | n=40 → n=80 | Report–proof consistency |
| 3 | swap_roi | Replace brain region | Cerebellum → Insula | Region axiom |
| 4 | drop_correction | Remove multiple-comparison correction | FDR/FWE markers deleted | Proof postcondition |
| 5 | drop_qc | Remove quality-control steps | Normality test, eddy current | QC axiom |
| 6 | fabricate_citation | Insert non-existent DOI | Fake DOI string | Axiom registry |
| 7 | modality_mismatch | Wrong tool for data modality | fMRI tool on DWI data | Tool contract axiom |
| 8 | wrong_stat_test | Misapply statistical test | Parametric on non-normal | Statistical axiom |
| 9 | pipeline_violation | Skip required preprocessing | Tensor fit before eddy | Dependency axiom |

Table S2: Per-Type Recall Comparison

| Injection Type | LeanNI Proof Kernel | LLM-as-Judge (DeepSeek-V3.2) |
|---|---|---|
| swap_p_value | 1.000 | 0.045 |
| swap_n | 1.000 | 1.000 |
| swap_roi | 1.000 | 0.927 |
| drop_correction | 1.000 | 0.155 |
| drop_qc | 1.000 | 0.000 |
| fabricate_citation | 1.000 | 0.982 |
| modality_mismatch | 1.000 | 1.000 |
| wrong_stat_test | 1.000 | 0.009 |
| pipeline_violation | 1.000 | 0.436 |

**Table S2.** Per-type recall for the two detectors (110 samples per type). The LLM judge exhibits a bimodal failure pattern: near-perfect recall on semantically salient violations (modality mismatch, sample-size manipulation, citation fabrication) but near-zero recall on subtle methodological omissions (QC removal, wrong statistical test, p-value falsification). The proof kernel achieves 100% recall uniformly across all types.

Table S3: Cumulative Ablation Study

| Layers | Configuration | Recall | FPR | ΔRecall |
|---|---|---|---|---|
| 1 | Kernel type-check | 0.444 | 0.0 | — |
| 2 | + Consistency check | 0.667 | 0.0 | +0.222 |
| 3 | + QC axiom | 0.778 | 0.0 | +0.111 |
| 4 | + Modality axiom | 0.889 | 0.0 | +0.111 |
| 5 | + Pipeline dependency | 0.889 | 0.0 | +0.000 |
| 6 | + Citation registry | 1.000 | 0.0 | +0.111 |
| 7 | + Statistical test | 1.000 | 0.0 | +0.000 |
| 8 | + Sentence audit | 1.000 | 0.0 | +0.000 |

**Table S3.** Cumulative ablation of detection layers. Layers are added incrementally; recall is measured over all 990 corrupted reports. The kernel type-checker alone covers 44.4% of injection types (4/9) through postcondition verification. Six layers suffice for 100% recall. FPR remains zero at all configurations, a property guaranteed by the axiom-grounded design. Layers 7–8 provide redundant coverage for robustness.

#### Supplementary Code: Proof Kernel (kernel.py)

The single entry point `check(pt, expected, ctx)` recursively type-checks a proof term against an expected proposition. The full source code is available at the project repository.

```
def check(pt: ProofTerm, expected: Prop, ctx: ProofContext) -> Prop:
    """Type-check a proof term against expected proposition.
    Returns the proven proposition or raises ProofError."""
    if ctx.depth > MAX_DEPTH:
        raise ProofError("depth limit exceeded")

    match pt:
        case AxiomRef(name=name, bindings=bindings):
            axiom = ctx.registry.get(name)
            if axiom is None:
```

```
        raise ProofError(f"unknown axiom: {name}")
    prop = instantiate(axiom.prop, bindings)
    for premise in axiom.premises:
        check(premise.proof, premise.prop, ctx.descend())
    if has_unbound_vars(prop):
        raise ProofError("unbound variables remain")
    return unify(prop, expected)

case ToolOutputRef(artifact_hash=h, postcondition=post):
    if h not in ctx.trusted_artifacts:
        raise ProofError(f"untrusted artifact: {h[:8]}...")
    return unify(post, expected)

case RuleApp(rule=rule, children=children):
    rule_fn = ctx.rules.get(rule)
    if rule_fn is None:
        raise ProofError(f"unknown rule: {rule}")
    child_props = [
        check(c, c.expected, ctx.descend())
        for c in children
    ]
    result = rule_fn(child_props)
    return unify(result, expected)

case Hole():
    raise ProofError("open proof obligation (Hole)")
```
